# Performance Evaluation of a Quantitative Metabolomics Workflow Incorporating Microchip Capillary Electrophoresis, Indexed Migration Time, and Single-Point External Calibration

**DOI:** 10.64898/2026.07.10.737294

**Authors:** J. Scott Mellors, Corinne Moss, Erin A. Redman, Christopher Shuford, James P. Campbell, J. Michael Ramsey, Joshua J. Coon, J. Will Thompson

**Author notes:** Corresponding Author: J. Scott Mellors.

## Abstract

Capillary electrophoresis-mass spectrometry (CE-MS) offers unique analytical advantages for polar metabolite profiling but has remained underutilized in metabolomics relative to liquid chromatography-MS (LC-MS), in part due to challenges in managing migration time drift during data analysis. Here we introduce the use of indexed migration time (iMT) for easily managing this aspect of CE-MS data for metabolomics. Migration time indexing using a panel of stable isotope-labeled (SIL) amino acid reference standards, stored as an iRT database in Skyline, outperformed both uncorrected migration time and relative migration time (RMT) correction across three independent analytical batches spanning 90 samples from four biological matrices. The indexed migration time approach achieved sub-1% relative standard deviation (RSD) in migration index across batches, compared to up to ~15% RSD for uncorrected migration times. Additionally, we evaluate the use of single-point external calibration in Skyline for the purposes of metabolite quantification from complex matrices in order to ease the burden of translational metabolite quantification from metabolomics using high-resolution mass spectrometry (HRMS). Single-point external calibration using a biological matrix-based calibrator was benchmarked against a 13-point linear calibration curve across a panel of amino acids; above 1 μM, greater than 95% of back-calculated concentrations fell within ±20% of multi-point calibration. Application of the complete workflow to plasma, serum, urine, and NIST Standard Reference Material (SRM)-1950 demonstrated low inter-batch variability by principal components analysis, broad metabolite coverage across 126 quantifiable analytes, and strong quantitative concordance (Deming slope = 0.862, pseudo-R^2^ = 0.994, n = 64 analytes) with an independent comprehensive reference dataset for NIST SRM-1950. Together, these results establish a practical mCE-HRMS metabolomics workflow that bridges targeted and discovery metabolomics paradigms and lays the groundwork for single-point external calibration as a powerful tool for translational metabolomics.

## Introduction

Capillary electrophoresis-mass spectrometry (CE-MS) has been established as a powerful analytical platform for metabolomics, particularly for the profiling of polar and charged metabolites that are poorly retained on conventional reversed-phase liquid chromatography (RPLC) columns.^1–3^The physicochemical basis for CE separations affords selectivity orthogonal to RPLC, resolving compounds on the basis of their charge-to-size ratio. Moreover, microchip CE separations (ZipChip®) are typically completed within minutes, and the inherently low volumetric flow rates (nL/min) facilitate highly efficient electrospray ionization, yielding excellent concentration sensitivity for polar analytes at the low-nM level (on-chip sample concentration).^4,5^ Despite these compelling characteristics, CE-MS remains dramatically underutilized relative to LC-MS in the metabolomics literature: a search of PubMed citations from 2020–2025 reveals approximately 100-fold fewer entries pairing “CE-MS” with “metabolomics” than those pairing “LC-MS” with “metabolomics,” indicating that adoption barriers extend well beyond analytical performance.

Perhaps the most prominent analytical challenges limiting adoption is migration time variability, which arises from fluctuations in capillary temperature and the variable ionic strength of biological matrices.^6^ This variability complicates compound identification and is handled poorly by most current metabolomics software, which was designed around the comparatively stable retention times of LC systems. Microchip CE (mCE) reduces separation path length and improves thermal dissipation, compressing the magnitude of these fluctuations and enabling faster analyses, but migration time drift remains a first-order concern compared to LC-MS.^7^ Nonetheless, given that there is no stationary phase in CE and migration order should theoretically behave much more like a fundamental physicochemical property than in LC, there is a compelling rationale to try and address this limitation. In theory, CE migration could then be standardized across laboratories and used as a fundamental metabolite identifier, much like collisional cross-section (CCS) has gained traction as an orthogonal metabolite identifier.^8–10^

Several mathematical frameworks have been proposed to normalize CE migration times. Relative migration time (RMT), in which migration times are normalized to a single reference compound spiked into each sample, is straightforward to implement and has been shown to improve metabolite annotation confidence.^11^ However, RMT corrections are practically accurate only in the immediate vicinity of the reference compound; deviations increase with distance from the reference peak, as migration time drift in CE is not linear across the electropherogram. Effective electrophoretic mobility (μ_eff_) addresses this shortcoming by converting migration times to a more absolute scale.^12^ A multi-laboratory ring trial demonstrated that conversion to μ_eff_ reduced inter-laboratory migration time variability from approximately 10.9% relative standard deviation (RSD) on the RMT scale to approximately 3.1% RSD, representing a significant improvement for compound annotation across platforms.^12^ Nevertheless, the μ_eff_ approach requires precise knowledge of experimental conditions, including capillary temperature and BGE viscosity, introducing sensitivity to temperature differences as small as 5 °C (equivalent to ~2% change in μ_eff_ per degree Celsius due to BGE viscosity changes).^12^ Furthermore, μ_eff_-based normalization is not compatible with transient isotachophoretic (tITP) focusing, a commonly used pre-concentration strategy for CE-MS that substantially improves sensitivity but decouples analyte migration from the model underlying μ_eff_.

Migration time indexing—converting observed migration times into a dimensionless index by interpolation against a defined panel of reference compounds—offers a conceptually elegant alternative that is both physically intuitive and practically straightforward. This approach is directly analogous to the Kováts retention index used routinely in gas chromatography for over six decades to report compound migration properties independent of specific experimental conditions,^13^ and to the indexed retention time (iRT) system developed for LC-MS/MS proteomics, in which a set of reference peptides defines a dimensionless scale that enables highly reproducible retention time prediction across laboratories and instrument configurations.^14^ In the present work, we adapt this migration time indexing strategy for mCE-MS metabolomics using a panel of stable isotope-labeled (SIL) amino acid standards as iMT markers within the Skyline software framework. An indexed migration time (iMT) approach has a key practical advantage over μ_eff_: it is agnostic to the absolute migration time model, and therefore compatible with tITP focusing, making it broadly applicable to state-of-the-art mCE-MS workflows.

Quantitative metabolomics has been equally challenged by an apparent dichotomy between targeted and untargeted approaches. Targeted methods using triple-quadrupole MS instruments with multi-point calibration curves deliver high precision and accuracy, but require extensive method development, offer no capacity for retrospective discovery, and are confounded by matrix effects if calibrators do not adequately reflect the biological sample.^15,16^ Discovery (non-targeted) metabolomics, typically implemented with high-resolution MS (HRMS) instruments, provides a hypothesis-agnostic survey of the metabolome but sacrifices inherent quantitative rigor and cross-study comparability.^17^ A recent push to bridge this divide through hybrid qual/quant or Simultaneous Quantitation and Discovery (SQUAD) HRMS workflows aims to maintain the coverage advantages of discovery metabolomics while delivering quantitative metabolite measurements analogous to targeted methods.^18,19^ Single-point external calibration has been proposed as a pragmatic tool within this framework: a single calibrator prepared at a known concentration is used to assign an analyte-to-internal-standard response factor for each metabolite, enabling quantitative reporting across a broad dynamic range with minimal calibration overhead.^20,21^ Prior work by van der Kloet et al. demonstrated that single-point calibration can substantially reduce systematic analytical error relative to simple semi-quantitative approaches, and Khamis et al. showed that single-point and multi-point calibration could agree within acceptable dynamic range limits.^20,21^ Nonetheless, the work of Khamis et al. examined a very limited number of analytes and stopped far short of an endorsement of single-point external calibration, perhaps limiting uptake in the field. Mid-level QC or study pool QC samples have frequently been utilized for inter-batch normalization for large targeted quantitative metabolomics cohort studies; this is effectively a type of single-point calibration which doesn’t utilize the quantitative value of the QC sample.^22–24^

Here we investigate these two established concepts—iMT indexing and single-point external calibration—in the specific context of microchip CE-HRMS metabolomics, using the ZipChip platform (Repligen Corporation) coupled to Orbitrap MS instruments (Thermo Fisher Scientific) and a Skyline-based quantitative analysis workflow.^25^ We demonstrate that (1) iMT, implemented using a set of SIL amino acid reference standards stored in a Skyline iRT database, substantially outperforms both uncorrected migration time and RMT for intra- and inter-batch compound identification; (2) a single-point external calibration approach yields quantitative performance within ±20% bias relative to multi-point linear calibration for concentrations spanning more than two orders of magnitude; and (3) the complete workflow enables reliable, quantitative metabolite profiling across diverse biological matrices, validated against an independent comprehensive reference dataset for NIST SRM-1950. More broadly, we hope to encourage use of migration time (and retention time) indexing and single-point external calibration as broadly useful and underutilized approaches for improving inter-laboratory translation of metabolomics measurements.

### Experimental Section

#### Reagents and Materials

Amino acid standard mixtures for system suitability testing were purchased from Promega Corporation (Madison, WI). A custom 36-component stable isotope-labeled (SIL) internal standard mixture was formulated by Cambridge Isotope Laboratories (Tewksbury, MA). Metabolite reference standards for library construction were purchased from IROA Technologies (Ann Arbor, MI) and Cerilliant Corporation (Round Rock, TX). NIST Standard Reference Material SRM-1950 was obtained from the National Institute of Standards and Technology. Biological specimens (human plasma, serum, and urine) were obtained from BioIVT (Westbury, NY). ZipChip consumables (HR chips) and background electrolyte (Peptides BGE) were purchased from Repligen Corporation. LC-MS grade methanol and ammonium acetate were purchased from Fisher Scientific (Waltham, MA). Vacuum filtration was performed with a 0.45 μm PTFE 96-well filter plate (Millipore, Burlington, MA). Calibration materials, quality controls, and blanks were created in batches (Move Analytical LLC) and stored at −80 °C prior to use.

#### Sample Preparation

Plasma, serum, and urine specimens were prepared using a methanolic protein precipitation protocol. Briefly, 20 μL of each biological sample, calibration standard or QC reagent was combined with 140 μL of extraction solvent (Methanol) containing the 36-component SIL internal standard mixture at 5 μM each. The samples were mixed for 15 minutes at room temperature followed by the addition of 40 μL focusing reagent (ammonium acetate in water) to support isotachophoretic focusing during CE injection. The mixture incubated at −20 °C for 10 minutes to complete protein precipitation and centrifuged at 4000 × g for 5 minutes at 4 °C to spin through the filter plate into the 96-well collection plate. During the biological matrix study, a study pool quality control (SPQC) sample was prepared by combining equal volumes of all extracted samples.

Calibration standards spanning 0.1 to 1000 μM were prepared by class-A dilution of a concentrated standard mixtures in 0.1 N HCl, followed by addition of extraction solvent containing internal standards. A single-point external calibration standard was prepared from spiked biological matrices, and value-assigned using an external reference laboratory (TMIC, Alberta, CA). All sample preparations were performed on ice.

### Microchip CE-MS Data Acquisition

Separations were performed on the ZipChip CE system (Repligen Corporation) using HR chips, Peptides BGE and an applied field strength of 500 V/cm. The mCE system was coupled to an Exploris 240 Orbitrap (Thermo Fisher Scientific, Waltham, MA). System suitability was assessed at the beginning of each analytical batch using an amino acid standard mixture (Promega) prepared in Peptides Diluent (908 Devices). Pass/fail criteria for system suitability were based on historical performance metrics from hundreds of analyses and included thresholds for migration time, peak resolution, and migration time. Samples were analyzed in randomized order within each batch, with SPQC injected every 8–10 samples and at the beginning and end of each batch. Two preparations of the single-point calibration standard were collected at the beginning and end of each batch, along with a single preparation of the Low QC (5 uM) and High QC (200 uM). For the 13-point calibration study, the calibration and QC samples were collected at the beginning and end of the batch.

### Indexed Migration Time (iMT) Database Construction

Indexed migration time was implemented using the iRT (indexed retention time) framework within Skyline (daily version, small molecule mode).^25,26^ A database of iRT values was constructed from the SIL internal standard mixture by analyzing diluted standards alongside biological samples and assigning each SIL compound a defined iRT value by linear interpolation between anchor standards spanning the full electrophoretic window. The resulting iRT-to-migration time calibration was applied within Skyline to convert all observed migration times to a dimensionless iMT score for each analytical batch. For comparison, relative migration time (RMT) was calculated by normalizing observed migration times to the migration time of ^13^C,^15^N-serine spiked into each injection. Raw migration time, RMT, and iMT were each evaluated as identifiers for seven representative amino acids across 90 samples spanning three independently prepared and analyzed batches and four biological matrices (plasma, serum, urine, and SPQC).

### Quantification – Multipoint Calibration

Quantification using multipoint calibration was performed in Skyline as previously described.^***25***^ For the calibration performance evaluation, a 13-point calibration curve (0.1–1000 μM) was analyzed, and the curated multi-point calibration (linear regression with 1/x weighting) was used as the reference; points outside 15% bias above the lower limit of quantification (LLOQ) and 20% at the LLOQ were excluded from multi-point curve fitting per standard method validation practice.^***27***^

### Quantification – Single point Calibration

Quantification using a single-point external standard approach within Skyline was performed as follows: (1) Analyte peak areas were normalized to the peak area of the corresponding SIL internal standard (or the nearest-migrating “surrogate” SIL compound when the labeled molecule was not available). (2) A calibration regression was defined between the Analyte/IS ratio for the reference material (“Single point calibrator”) and zero. In Skyline, the “Concentration Multiplier” field was used to set the value for the reference material for each analyte and Linear Through Zero was selected as the regression method. (3) Concentrations were assigned for unknowns by plugging the Analyte/IS ratio measured for each sample in step 1, into the regression established in step 2. For the comparison to multi-point calibration, quantitative performance of single-point calibration was evaluated by back-calculating concentrations for all remaining calibration points and comparing to the accepted multi-point concentrations. Bias was calculated as (C_calc_ − C_ref_) / C_ref_ × 100%, and the percentage of datapoints within ±20% bias was reported.

### Principal Components Analysis and Metabolite Coverage

For the multi-matrix performance evaluation, principal components analysis (PCA) was performed in Python (v3.10) using scikit-learn on log-transformed, mean-centered, and unit-variance-scaled data from analytes with less than 50% coefficient of variation (CV) in the SPQC and less than 50% missing data across all measurements (126 metabolites meeting both criteria). Metabolite coverage was assessed by tabulating the number of analytes with a valid measurement (non-missing, within the calibrated range) in each sample type, stratified by metabolite class.

Quantitative values for NIST SRM-1950 were compared to published reference values from Mandal et al. (2025),^28^ which provided a comprehensive, quantitative analysis of SRM-1950 across 64 metabolites in common with our CE-MS target library. Quantitative agreement was assessed by Deming regression.

## Results and Discussion

### Overview of the mCE-HRMS Metabolomics Workflow

The end-to-end workflow introduced in this study comprises four sequential phases: (1) sample preparation, (2) mCE-HRMS data acquisition, (3) peak assignment and targeted quantification in Skyline, and (4) quality assessment and statistical analysis (Figure 1). Biological specimens (plasma, serum, urine, and cell culture media) are extracted by methanolic protein precipitation in the presence of 36 SIL internal standards and ammonium acetate, which serves a dual role as a BGE-compatible additive and as the leading electrolyte for tITP stacking during CE injection. The SIL standards serve both as iMT indexing markers and as paired internal standards for analyte/IS ratio-based quantification. Data-dependent acquisition on Orbitrap HRMS instruments captures both MS1 accurate masses and MS2 fragmentation spectra, enabling both targeted quantification and retrospective non-targeted analysis from the same data files. A pooled single-point calibration standard, prepared in the same extraction matrix as the study samples, is analyzed alongside unknowns to enable absolute concentration assignment. All steps from raw data import through quantitative reporting are automated within a custom Skyline template workflow, with optional manual review of flagged peak assignments.

**Figure 1.**
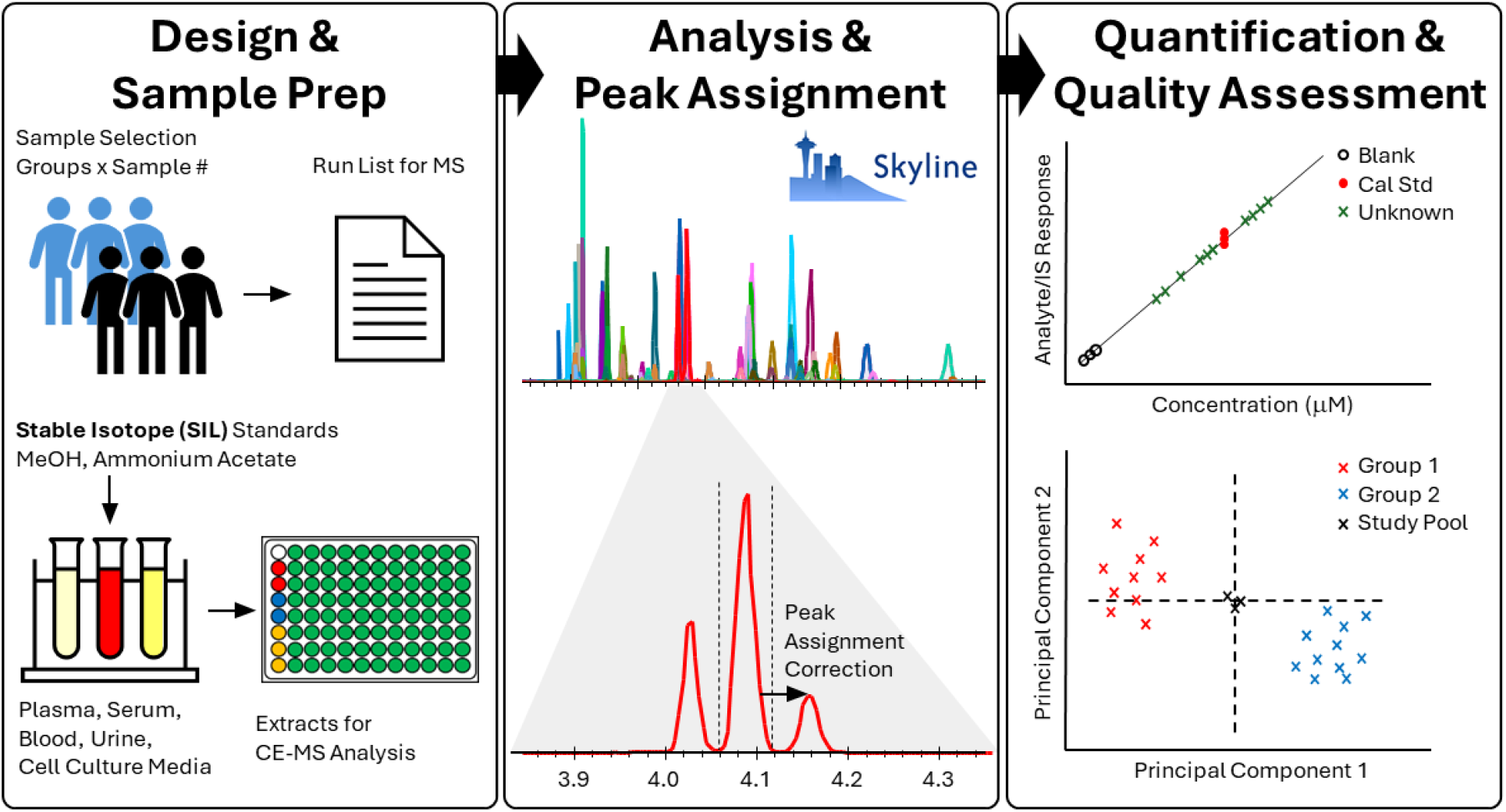
Schematic overview of the microchip CE-MS metabolomics workflow. *Left:* Sample preparation involves selection of study specimens (plasma, serum, urine, cell culture media), extraction by methanol protein precipitation including 36 stable isotope-labeled (SIL) internal standards and ammonium acetate, and vacuum filtration to remove protein precipitate. *Center:* CE-HRMS data are acquired using ZipChip® coupled to an Orbitrap mass spectrometer. Data analysis is performed in a Skyline-based pipeline: raw data are imported and peaks are assigned using the indexed migration time (iMT) framework. Right: Analyte/IS response ratios are converted to absolute concentrations using a regression between a single-point external calibrator and zero, and data quality and statistical assessments (including PCA) are performed.

### Indexed Migration Time Corrects for Migration Time Variability

Migration time variability is a primary practical barrier to the implementation of CE-MS metabolomics.^6,11^ Figure 2A illustrates the magnitude of this challenge: extracted ion electropherograms for eight amino acids from a QC sample analyzed in two different sample batches show a clear shift in absolute migration times. Plotting the same data against iMT (Figure 2B) eliminates this batch-dependent offset, aligning peaks from both batches across the full separation window.

**Figure 2.**
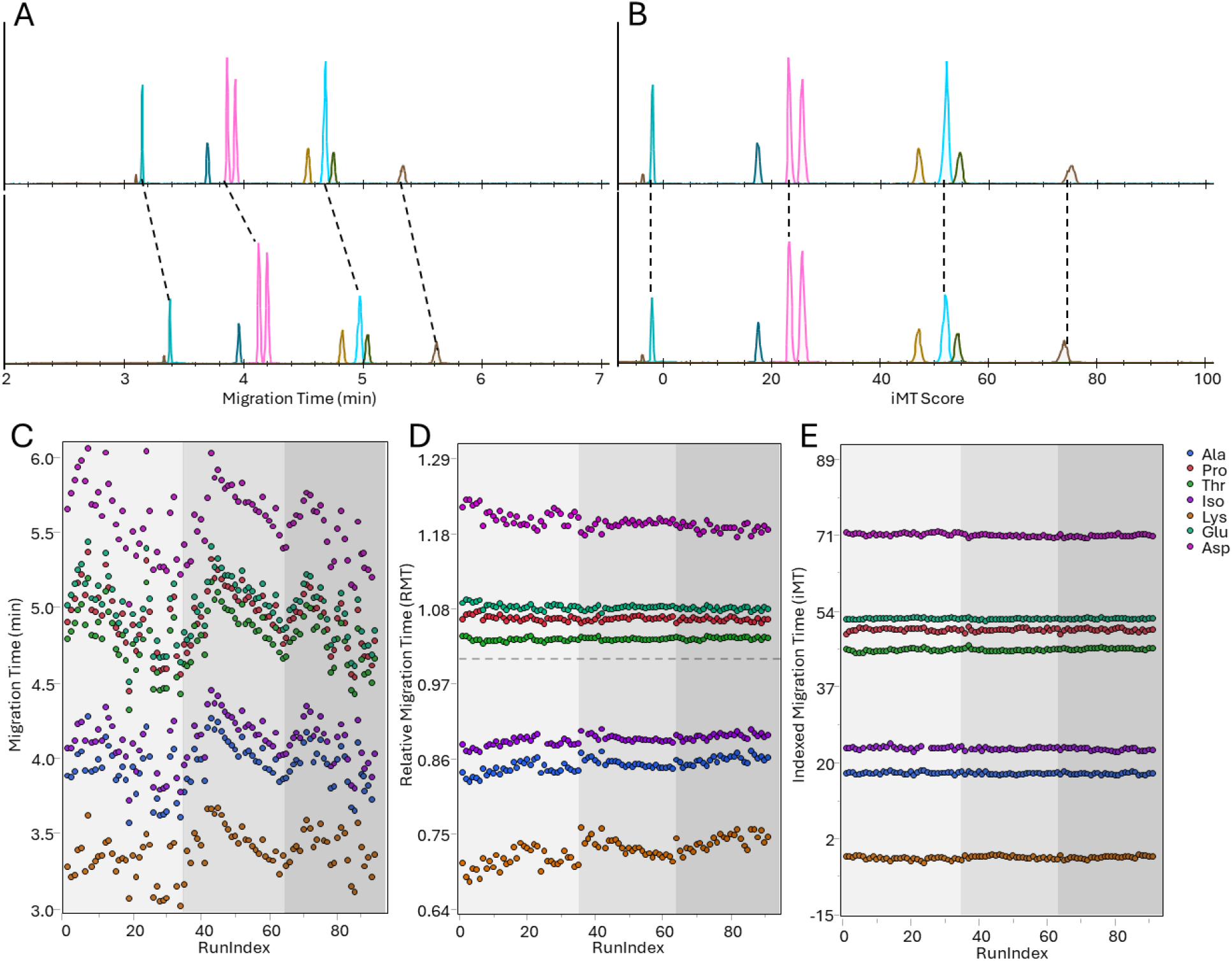
Indexed migration time (iMT) corrects for CE migration time drift across analytical batches. (A) Extracted ion electropherograms for eight amino acids from a 200 μM QC sample analyzed in batch 1 (top) and batch 3 (bottom), illustrating the shift in absolute migration times across batches. (B) The same samples plotted on the iMT scale, showing alignment of peaks from both batches. (C) Raw migration time plotted for seven representative amino acids (see legend) extracted from plasma, serum, urine, and QC matrices across 90 samples and three batches; shaded regions denote individual batches. (D) Relative migration time (RMT), normalized to ^13^C,^15^N-serine (dashed line at RMT = 1.0), showing effective correction near the reference compound but increasing deviation for more distant analytes. (E) Indexed migration time (iMT) for the same dataset, demonstrating near-uniform correction across the full migration window and consistent precision below 1% RSD for all analytes. The same raw migration time axis range is displayed in panels C–E to facilitate comparison.

To quantify the performance of migration time correction approaches across the full dataset, we tracked the migration time (raw, RMT, and iMT) for seven representative amino acids across 90 samples from three batches and four matrices. Raw migration times showed variability of up to007E15% RSD across batches (Figure 2C), primarily reflecting inter-batch BGE ionic strength differences and similar to previously reported results for CE.^12^ The previously-described RMT correction (Figure 2D), using ^13^C,^15^N-serine as the normalizing compound, substantially reduced variability for compounds migrating near serine (average migration time 007E4.7 min) but was less effective for compounds migrating considerably earlier or later, consistent with the known limitation that RMT corrections are only accurate in the immediate vicinity of the reference standard.^11^ In contrast, iMT (Figure 2E), calculated by linear interpolation with a subset of 17 internal standards spanning the full electrophoretic window, reduced migration time variability to below 007E1% RSD for all analytes regardless of their position in the electropherogram—a level of precision analogous to collision cross-section (CCS) values reported for ion mobility spectrometry. The absolute performance advantage of iMT over RMT across the full electropherogram is further visualized in Supplementary Figure 1, which plots RSD versus average migration time for all 22 canonical amino acids; RMT performs comparably to iMT only in the immediate vicinity of the RMT reference standard (serine), and iMT consistently achieves lower variability elsewhere.

**Supplementary Figure 1.**
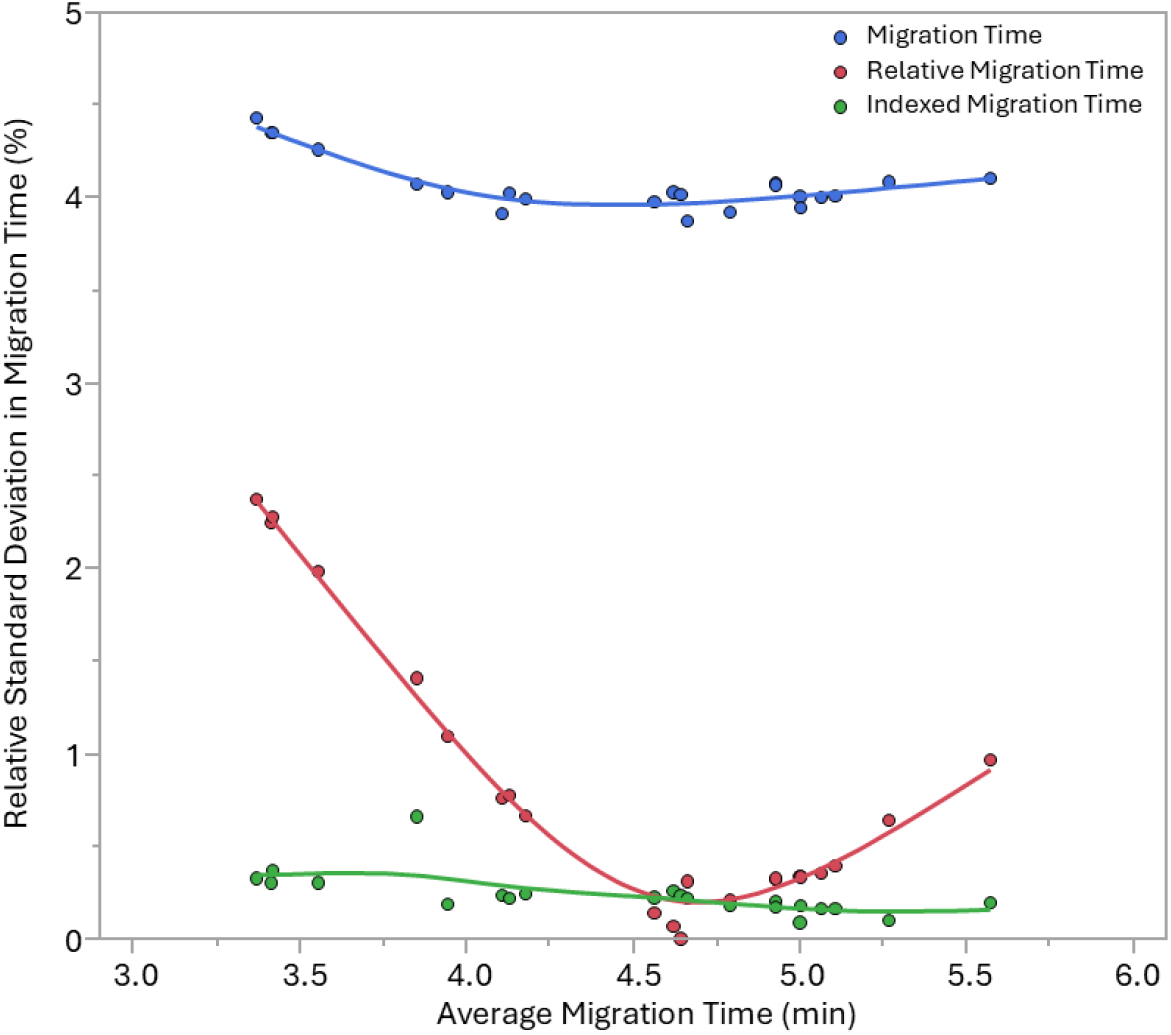
Relative standard deviation (% RSD) in migration time versus average migration time for 22 canonical amino acids across three batches of Experiment 1 (n = 90 samples), comparing uncorrected migration time (gray), RMT (orange), and iMT (blue). RMT achieves performance comparable to iMT only in the immediate vicinity of the RMT reference standard (^13^C,^15^N-serine, average migration time 007E4.7 min, denoted by dashed vertical line). iMT consistently achieves RSD below 1% across the full migration window.

### Remaining issues with peak assignment after iMT

Among the most time-consuming challenges in targeted metabolomics data analysis is the systematic misassignment of peaks for closely co-eluting compounds with similar or the same mass, which often require laborious manual curation.^19^ This problem is equally apparent in iMT-aligned mCE-MS data and LC-MS data. An example is shown in Figure 3 from the CE-MS workflow described in this manuscript: isoleucine (Ile), leucine (Leu), and alloisoleucine (allo-Ile) yield multiple peaks in the extracted ion electropherogram that must be assigned individually. Figure 3A shows three distinct peaks are resolved in the [M+H]^+^ electropherogram for one run, along with the indexed migration time (iMT) score predicted for isoleucine. Note that the time prediction is within 2 seconds (0.03 min) of the peak apex, but nonetheless imperfect. The default Skyline peak assignment, which selects the best candidate peak for each target independently, assigns the same (most abundant) peak to all three analytes in nearly all samples (Figure 3B), failing to leverage the knowledge that each target has a distinct predicted iMT and that two different targets cannot occupy the same peak. Green indicates correct assignments and red indicates incorrect assignments.

**Figure 3.**
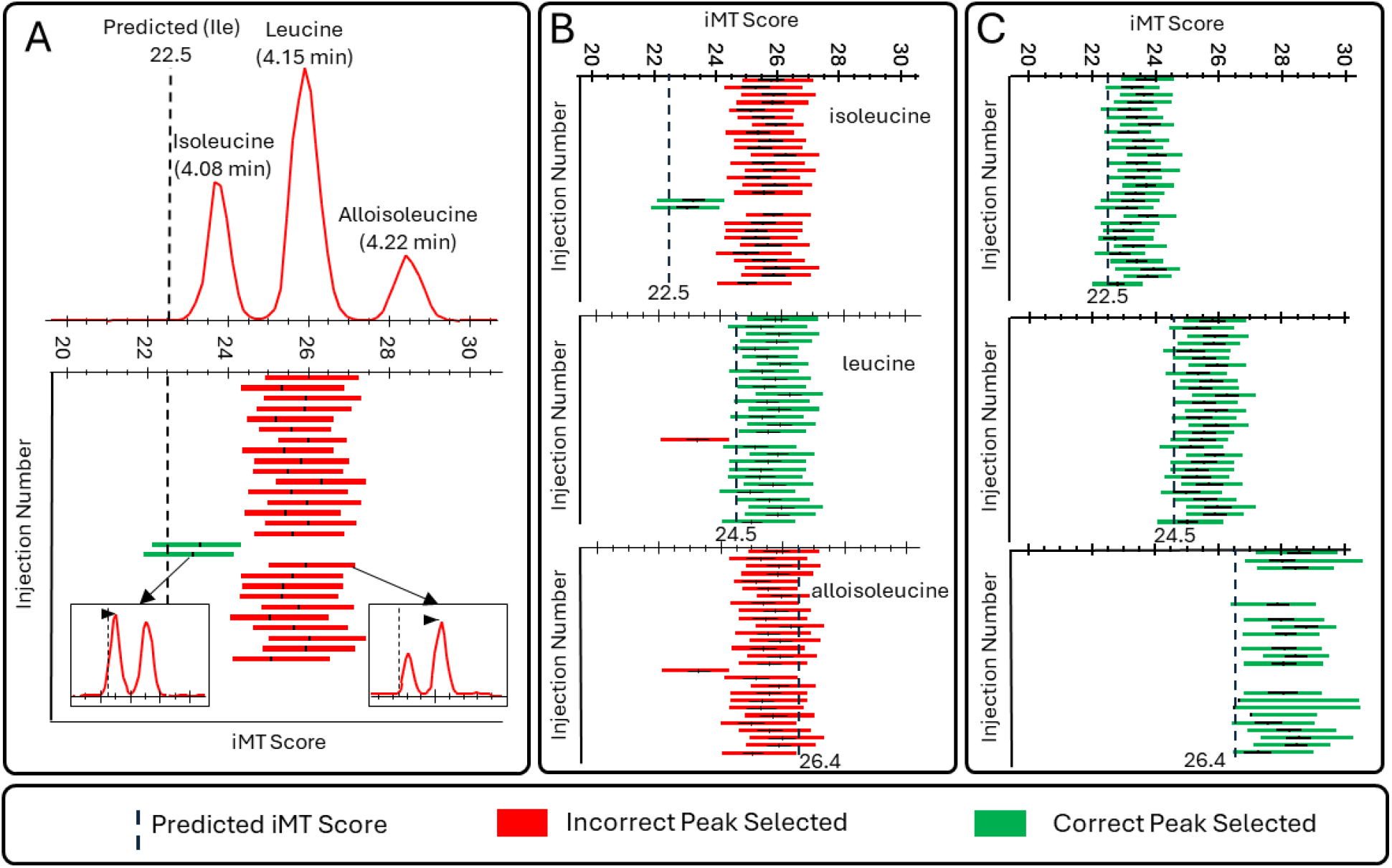
Peak assignment problem for closely eluting isobaric metabolites. (A) Upper panel: mCE-MS separation of isoleucine (Ile), leucine (Leu), and alloisoleucine (allo-Ile) in a biological sample spiked with additional allo-Ile, showing three resolved peaks at the shared [M+H]^+^ m/z. Lower panel: default Skyline peak assignment for isoleucine (predicted iMT = 22.5) across the analytical batch, where only two samples (green) were correctly assigned; most samples were assigned the most abundant peak (red). (B) Default Skyline assignments for Ile (top), Leu (middle), and allo-Ile (bottom) across the batch, with green indicating correct assignment and red indicating incorrect assignment; the same (most abundant) peak was assigned to all three analytes in nearly all samples. (C) Corrected assignments.

Figure 3C shows the corrected assignments for Ile, Leu, and allo-Ile along with their predicted iMT Score (vertical dashes), after manually correcting the peak assignment. Algorithmic approaches for automatically correcting these peak assignments is beyond the scope of this manuscript; the purpose of including it here is to say that while the iMT approach effectively brings CE-MS metabolomics on-par with LC-MS from a peak-time reproducibility standpoint, it does not completely address all peak assignment issues for correctly integrating closely eluting isobaric peaks.

### Single-Point External Calibration (SPEC) Provides Quantitative Performance Comparable to Multi-Point Calibration over a Wide Dynamic Range

Targeted quantitative metabolomics conventionally employs multi-point calibration curves (typically 7–13 points spanning 3–4 orders of magnitude) to characterize the linear dynamic range and assign absolute concentrations.^27,29^ While rigorous, this approach requires substantial resources, including reagent costs for authentic standards at all calibration levels and instrument time to analyze the full curve for each batch. In contrast, untargeted studies typically only employ relative quantification, effectively creating isolated islands of information and limiting translation between laboratories and experiments. Our hypothesis is that for modern instruments with high linear dynamic range, a single-point external calibration (SPEC) could serve as a more efficient alternative that is particularly attractive for large-scale discovery-to-quantification workflows where the full range of analyte concentrations is not known a priori.^20,21^

To evaluate this approach in the context of mCE-HRMS metabolomics, we compared SPEC to a 13-point multi-point linear calibration (with 1/x weighting) for a panel of amino acids spanning 0.1– 1000 μM. Figure 4A shows a representative multi-point calibration curve for valine (R^2^ = 0.9962, slope = 0.9313 with 1/x weighting), with the LLOQ-based curation excluding the 0.1 and 0.25 μM points for this analyte. Figures 4B and 4C show the analogous single-point calibration curves using the 1 μM and 100 μM calibrators, respectively, with all other concentrations treated as quality controls (i.e. left out of the regression but with their calculated concentrations reported on the plot).

**Figure 4.**
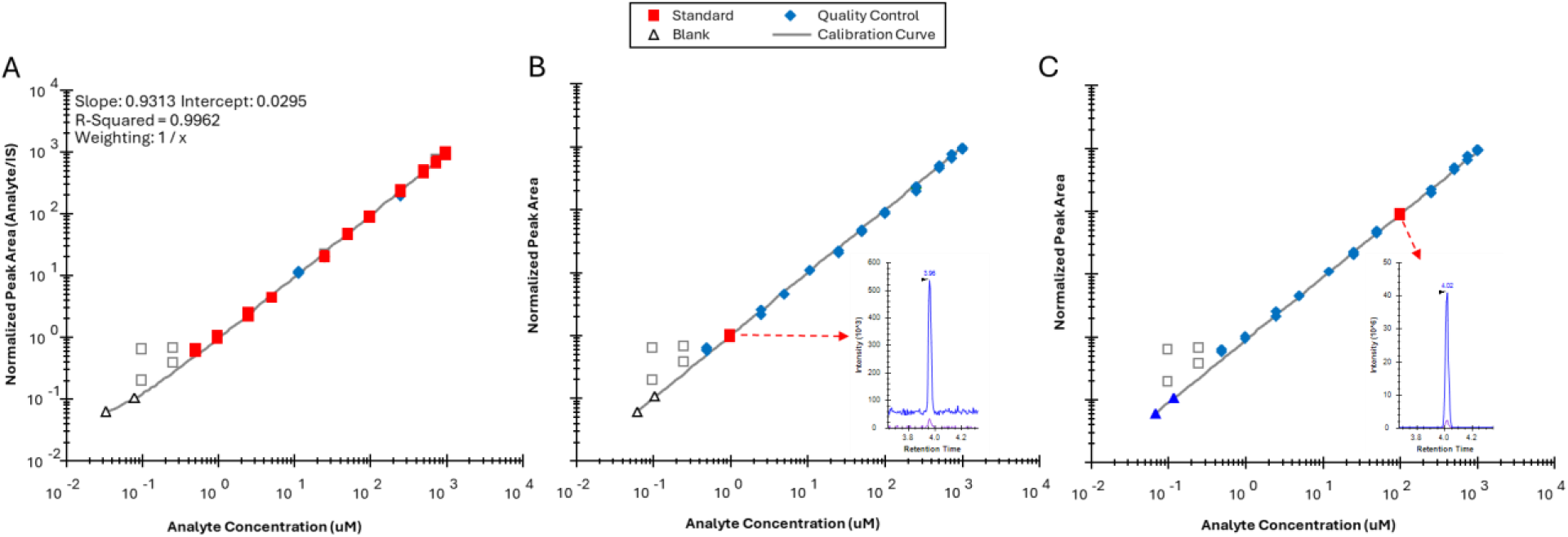
Comparison of multi-point and single-point external calibration strategies for mCE-HRMS quantification. (A) Thirteen-point calibration curve for valine using linear regression with 1/x weighting (R^2^ = 0.9962, slope = 0.9313); points at 0.1 and 0.25 μM were excluded following LLOQ-based curation (filled symbols = included; open symbols = excluded). Single-point calibration using the 1 μM calibrator (fit through zero); remaining concentrations are treated as quality controls. (C) Single-point calibration using the 100 μM calibrator. Insets in (A) and (B) show the extracted ion electropherogram for valine at the calibration concentration used for single-point fitting.

Across the panel of amino acids, after curation of the multi-point calibration curves, there were 590 data points within ±20% bias in the multi-point analysis. Using the 100 μM single-point calibrator (Figure 4C), back-calculated concentrations for all other calibration levels were compared to the multi-point reference: at 1 μM and above (spanning a 1000-fold range around the calibrator), greater than 95% of measurements (514 of 541) maintained better than ±20% bias (Supplementary Figure 2). Performance deteriorated at 0.5 μM (34% of measurements within 20% bias) and was unreliable below 0.5 μM, a region where both the multi-point and single-point approaches performed poorly.

It is important to note that all of the points in the multi-point approach in Supplementary Figure 2 (top) were included in the regression itself (Back-Fit Accuracy). However, only a single point in the lower panel of the figure was included in the regression, representing an independent assessment of accuracy from the other points. Therefore, this experiment may underrepresent the true relative performance of SPEC compared to traditional multi-point calibration.

**Supplementary Figure 2.**
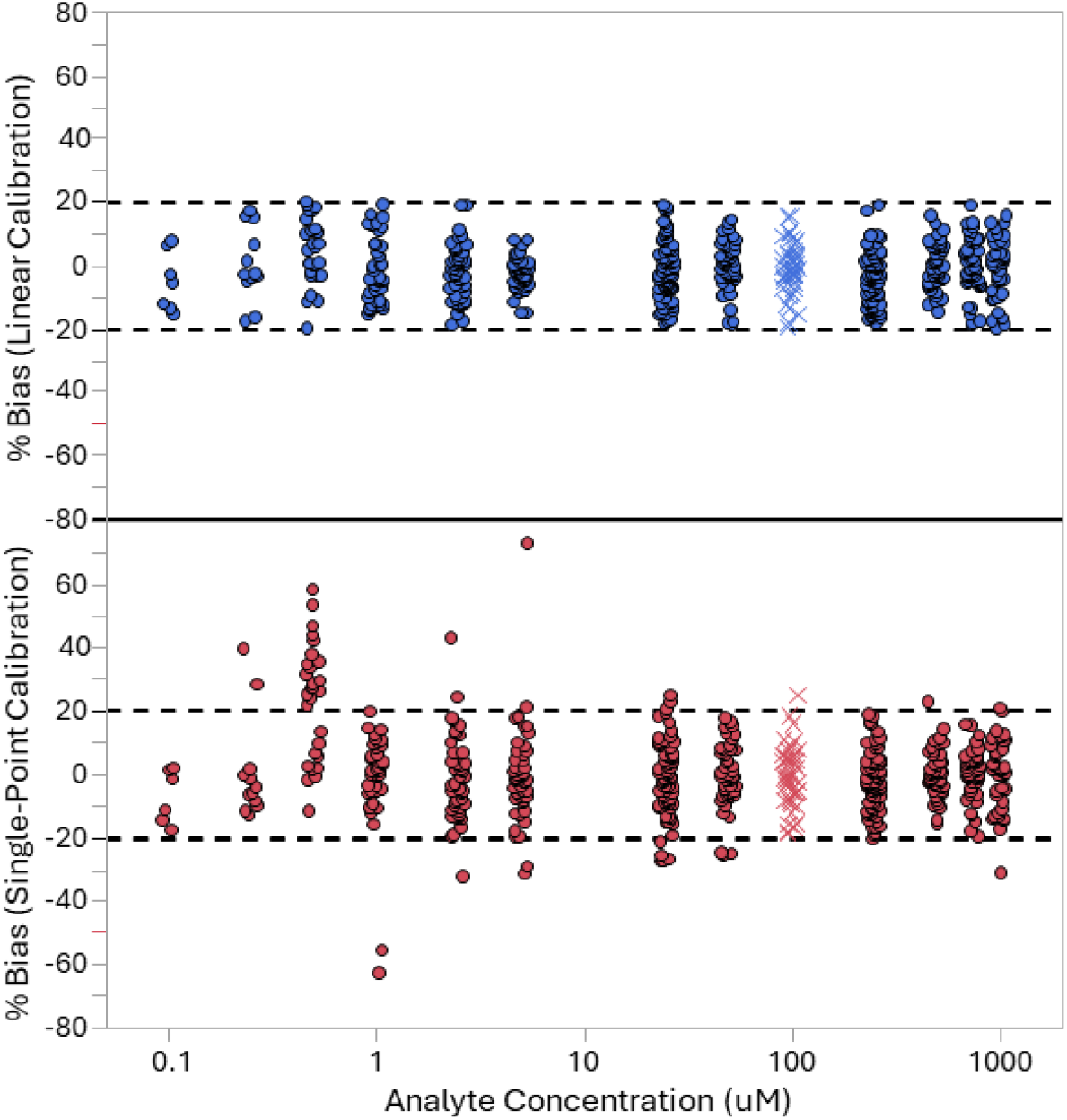
Comparison of percent bias distributions for 13-point linear calibration (blue) versus single-point calibration at 100 μM (red) across the amino acid panel. Red hashes denote the single-point calibrator concentration (100 μM). At concentrations of 1 μM and above, 95% of single-point back-calculated concentrations (514/541 measurements) fell within ±20% bias of the multi-point reference, confirming quantitative equivalence over the useful dynamic range. Bias for both methods increases markedly below 0.5 μM, which represents the practical LLOQ for this analyte panel on the mCE-HRMS platform.

### Requirements and Limitations of Single-Point External Calibration

The results demonstrate that within the linear dynamic range of the Orbitrap detector, SPEC using a calibrator within 007E10-fold to 100-fold of the unknowns can deliver quantitative accuracy that is practically indistinguishable from multi-point calibration—at a fraction of the analytical overhead. A few important caveats exist that could limit practical application of SPEC in some cases. First, the approach requires that there be enough signal to estimate the response-per-mol for that analyte; i.e., the peak must be in the linear response range of the detector. This will limit the utility of SPEC for very low abundance metabolites. Second, a limitation of the approach that must be acknowledged is that the experiment does not directly define lower and upper limits of quantification in each batch. Similar to multi-point calibration, quality control (QC) samples at known low and high values can be used to verify performance in each batch across a wide metabolite dynamic range.

### Workflow Performance across Biological Matrices

To assess the performance of the complete iMT-indexed, single-point calibration workflow in a realistic multi-sample study setting, we applied it to three independent analytical batches that included NIST SRM-1950 (n ≥3 preparations per batch), single-donor human plasma, serum, and urine specimens, and a SPQC sample prepared from equal contributions of all specimen types.

Figure 5A shows a PCA of all samples using the 126 analytes meeting data quality thresholds (≤50% CV in SPQC, ≤50% missing data across all measurements). Samples cluster clearly by matrix type along the first two principal components with no apparent inter-batch clustering, indicating that the iMT-based alignment and single-point calibration approach effectively normalizes between-batch analytical variability. The SPQC cluster falls at the intersection of PC1 and PC2, as expected for a pooled average of all samples, confirming that the SPQC accurately represents the overall study composition and is measured without systematic bias.

**Figure 5.**
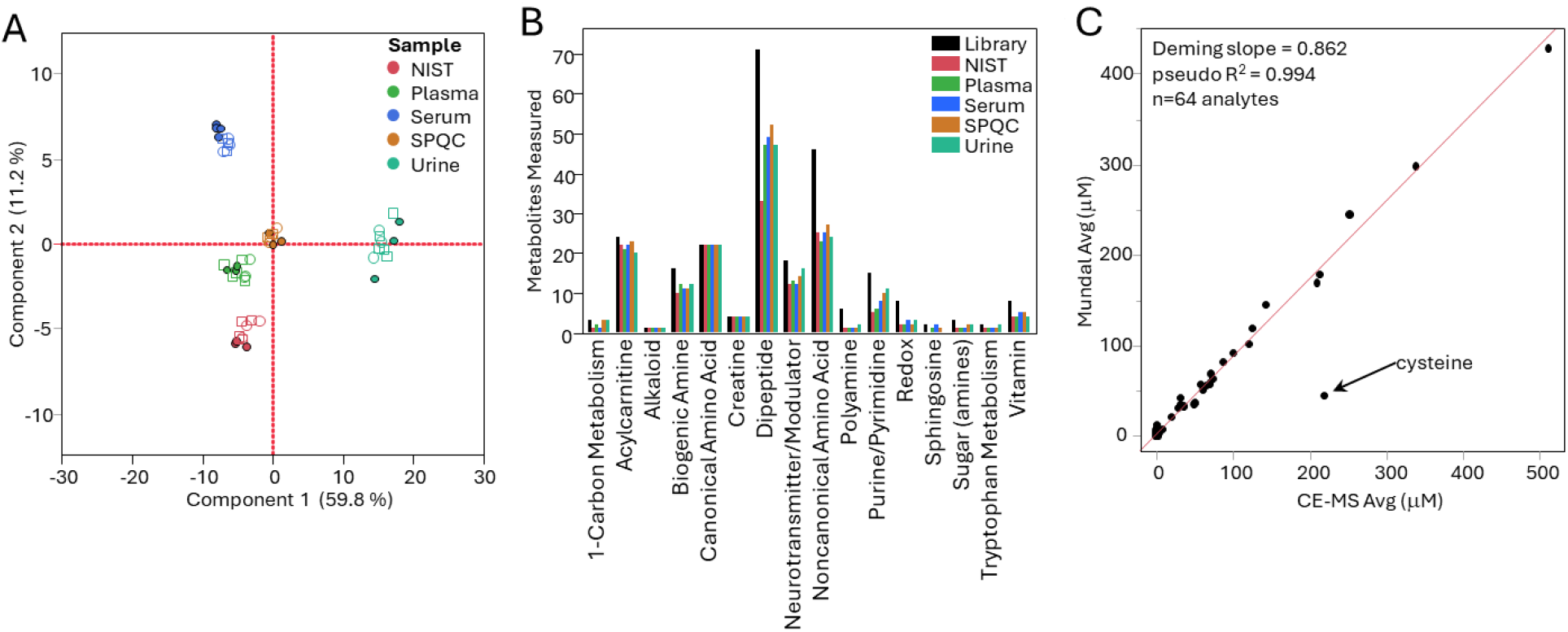
Performance of the end-to-end mCE-HRMS workflow across multiple biological matrices. (A) Principal components analysis (PCA) using 126 analytes with ≤50% CV in SPQC and ≤50% missing data across all measurements. Samples cluster by matrix type with no apparent inter-batch clustering; the SPQC (gray) plots at the centroid of all samples (PC1: 59.8%, PC2: 11.2%). (B) Metabolite coverage by sample type and metabolite class; the Library bar represents the full 325-compound target list. SPQC shows the broadest coverage due to contributions from all matrices. (C)Quantitative comparison of NIST SRM-1950 concentrations measured by the mCE-HRMS single-point calibration workflow versus published reference values from Mandal et al. (2025) for 64 metabolites in common (Deming slope = 0.862, pseudo-R^2^ = 0.994). Cysteine (excluded from regression, asterisk) is elevated in the CE-MS data due to the addition of a reducing agent during sample preparation, which converts disulfide-bound cysteine to the free form.

Figure 5B summarizes metabolite coverage across matrices and metabolite classes. The 325-compound CE-MS target library spans 16 metabolite classes including canonical and non-canonical amino acids, acylcarnitines, biogenic amines, purines and pyrimidines, dipeptides, 1-carbon metabolism intermediates, polyamines, and vitamins. SPQC, because it includes contributions from all matrices, shows the highest coverage across categories (as expected). Urine exhibited the lowest coverage. The breadth of coverage across diverse chemical classes underscores the utility of CE-MS with its tITP-enhanced sensitivity for profiling the polar metabolome without the need for chemical derivatization.

To benchmark the quantitative accuracy of the single-point calibration approach against orthogonal data, we compared CE-MS concentrations for NIST SRM-1950 to published reference values from Mandal et al. (2025),^28^ who performed a comprehensive quantitative characterization of SRM-1950 using an independent platform. Of the 64 metabolites in common between the two datasets, Deming regression yielded a slope of 0.862 and a pseudo-R^2^ of 0.994 (Figure 5C), indicating excellent linearity and good quantitative agreement across more than two orders of magnitude in concentration. The slight systematic deviation from unity slope may reflect calibration differences between platforms, matrix effects in the SRM-1950 material, or differences in the specific isoforms measured by each assay. One notable outlier was cysteine, for which our CE-MS values were higher than the Mandal reference; this is consistent with the addition of a reducing agent to our sample preparation, which would convert cystine and mixed disulfides to free cysteine, increasing the apparent cysteine concentration. This example highlights the importance of considering pre-analytical factors when comparing metabolite concentrations across platforms and laboratories.

## Conclusions

We have demonstrated that two established analytical concepts, migration time indexing (iMT) and single-point external calibration (SPEC), can be productively combined within a microchip CE-HRMS metabolomics workflow to address the most practically significant limitations of CE-MS for metabolomics at scale. iMT, implemented using SIL internal standards as indexing markers within the Skyline iRT framework, achieves cross-batch migration time precision below 1% RSD across diverse biological matrices, substantially outperforming both uncorrected migration time and RMT approaches, particularly for analytes distant from the RMT reference compound. Single-point external calibration shows promise to achieve quantitative accuracy within ±20% bias for analytes with concentrations in the linear response range of the MS detector. We therefore believe that value-assigned complex (or spiked) biological matrices can provide an efficient, discovery-compatible alternative to traditional multi-point calibration. Application of the full workflow across plasma, serum, urine, and NIST SRM-1950 demonstrated low inter-batch variability, broad metabolite coverage across 16 compound classes, and strong quantitative concordance with an independent reference dataset. Together, these results provide a rigorous quantitative foundation for mCE-HRMS as a practical, end-to-end metabolomics platform suitable for multi-batch and multi-matrix clinical and biological studies.

